# An evaluation of screening methods for the detection of extended-spectrum beta-lactamase-producing *Escherichia coli* and *Klebsiella pneumoniae* in environmental samples from healthcare settings

**DOI:** 10.1101/2025.08.12.669680

**Authors:** Esther Picton-Barlow, Claudia McKeown, Sally Forrest, Sarah Gallichan, Maria Moore, Nicholas A. Feasey, Joseph M. Lewis, Fabrice E. Graf

**Affiliations:** Department of Clinical Sciences, Liverpool School of Tropical Medicine, Liverpool, United Kingdom; School of Medicine, University of St Andrews, St Andrews, United Kingdom; Tropical Infectious Diseases Unit, Royal Liverpool University Hospital, Liverpool, United Kingdom

**Keywords:** Antimicrobial resistance, Surveillance, Culture methods, Infection control, Healthcare-associated infections, Pre-enrichment, Microbiology

## Abstract

Drug-resistant infections with extended-spectrum beta-lactamase-producing Enterobacterales (ESBL-E) are increasingly problematic, especially in healthcare settings. ESBL-E infections lead to a worse treatment outcome and often require escalating antibiotic treatment to reserve antibiotics such as carbapenems. Understanding how these bacteria are transmitted within healthcare settings is complex and not well understood, but it is a requirement for effective infection prevention and control strategies. To what extent environmental ESBL-E reservoirs contribute to transmission is unclear. To accurately capture these reservoirs, laboratory processing of environmental samples needs to be optimised to reliably detect ESBL-E, which is challenging because they can be scarce and part of complex microbial communities.

Here, we assessed screening methods for the detection of ESBL-producing *Escherichia coli* (ESBL-Ec) and *Klebsiella pneumoniae* (ESBL-Kp) from environmental swabs taken from healthcare settings, testing different swab types, pre-enrichment conditions and selective agars.

We show that a pre-enrichment of 18 hours significantly increased the recovery of 3GC-resistant gram-negatives. The choice of selective agar impacted the number of ESBL-Ec and ESBL-Kp detected. This also affected the number of samples requiring additional species confirmation, which is costly and time-consuming. For our use case and by considering additional factors such as cost and practical aspects, Membrane Lactose Glucuronide Agar supplemented with cefotaxime performed best for the combined detection of ESBL-Ec and ESBL-Kp.

Our work will guide environmental surveillance of ESBL-E by providing optimised methods for laboratory processing of environmental samples from healthcare settings.

## Introduction

Infections with extended-spectrum beta-lactamase-producing Enterobacterales (ESBL-E) are associated with high mortality, morbidity and healthcare costs (1–3). These infections are challenging to treat and often necessitate the use of carbapenems or other Watch and Reserve antibiotics. To lessen burden of ESBL-E infections and reduce reliance on reserve antimicrobial therapies, it is essential to reduce the spread of ESBL-E. To interrupt their spread requires a better understanding of transmission routes to inform targeted infection prevention and control measures.

Healthcare settings provide an increased opportunity for transmission of drug-resistant bacteria between people due to a combination of drivers, including increased antibiotic use, breakdown of hygiene, increased contact with healthcare workers, invasive procedures and devices, and comorbidities, including compromised immune systems (4). Additionally, the healthcare environment, e.g. high-touch surfaces, shared bathrooms, etc, presents an extra point of contact that can become contaminated with drug-resistant bacteria and serve as a reservoir of infection (5–7). However, the importance, routes and directionality of these reservoirs are not yet well understood.

Detection of drug-resistant bacteria in the healthcare environment is challenging because the pathogen of interest may be present at low levels and part of a complex microbial matrix, such as dry surface biofilms and hydrated biofilms (8,9). In general, the workflow to detect these bacteria usually starts with swabbing the environment, followed by pre-enrichment in nutrient broth, screening using selective agar plates, and then species confirmation and sometimes additional test e.g. antibiotic susceptibility testing. Additional characterisation, such as whole genome sequencing (WGS), are rarely performed (10) to determine genetic relatedness of bacteria, which allows inference of transmission, and has been successfully used in outbreak investigations (5,11). There is, however, a lack of consensus on optimal approaches to microbiological surveillance of healthcare environments.

Pre-enrichment of swabs in nutrient broth can promote the growth of organisms before plating onto agar, but is sporadically applied in environmental sample processing, and there is no consensus for the incubation duration and broth type (10). One commonly used broth for ESBL-E enrichment is Tryptic Soy broth (TSB), a high-nutrient medium that has substantially improved detection of ESBL-producing bacteria in clinical samples compared to direct plating (12,13). Buffered Peptone Water (BPW) is a recovery medium commonly used to aid recovery of bacteria from environmental, food and wastewater samples (14) .

Similarly, the selective agar with the highest bacterial recovery is unclear and likely context specific. The high species diversity in the environment makes it challenging to find an agar with selective properties that remains sensitive to the full range of potential pathogens of interest. Plating onto MacConkey agar supplemented with a 3GC (e.g. cefotaxime) is a popular and globally used technique for detecting ESBL-E (15,16). Chromogenic agar, such as CHROMagar ESBL, is increasingly favoured because of its ability to distinguish between different ESBL-producing species using colour. However, chromogenic agar has been reported to be markedly less specific than supplemented MacConkey, especially in settings where ESBL-E are commonplace (16). Alternatively, species-specific agar can be supplemented with a 3GC to screen for particular ESBL-E. This includes *E. coli*-differential media, such as Membrane Lactose Glucuronide Agar (MLGA) (17) and *Klebsiella*-differential media such as Simmons Citrate with inositol (SCAI) (18).

In this study, we evaluate screening methods most suitable for environmental surveillance in healthcare settings for the detection of ESBL-producing *E. coli* (ESBL-Ec) and *Klebsiella pneumoniae* (ESBL-Kp). We compare the recovery of ESBL-Ec and ESBL-Kp from swab samples taken from healthcare environments (sinks, showers, toilets and high-touch surfaces) when pre-enriched in two nutrient broths (BPW and TSB) at 4 and 18 hours. We also evaluate four different agars (CHROMagar ESBL and cefotaxime-supplemented MLGA, MacConkey and SCAI) on their suitability for ESBL-Ec and ESBL-Kp screening, assessing their detection capabilities. Lastly, we consider the practical and financial considerations involved with each method.

## Materials and Methods

### Sample collection

We performed a series of experiments to compare different swabs, different pre-enrichment conditions, and selective agar. No sample size calculation was carried out; the number of samples was a convenience sample. Environmental samples were taken from shower heads, shower drains, handwash sinks, toilets, and high-touch surfaces (Table 1) in hospital wards and care facilities across Liverpool, as part of the Tracking AMR across Care Settings (TRACS-Liverpool) study (19) . Surfaces were swabbed for 20 seconds, rotating the swab throughout. Initially, sites were sampled using a polyester swab (TS/19-G250, Technical Service Consultants) and a foam swab (ENVSWB100, Neogen) to evaluate swab performance. Subsequently, only polyester swabs were used, moistened with sterile PBS directly before use. Our swabbing approach was adapted from (20,21). Swabs were transported from the sampling site to the laboratory at room temperature within 7 hours and processed within 2 hours of reaching the laboratory.

**Table 1:** Number of collection sites within hospitals and care facilities that were sampled for each method evaluation in this study.

|  | Swab comparison | Pre-enrichment | Agar |
| --- | --- | --- | --- |
| Sink | 14 | 35 | 49 |
| Toilet | 7 | 11 | 10 |
| Shower drain | 8 | 16 | 20 |
| Shower head | 0 | 4 | 1 |
| High-touch surfaces | 1 | 16 | 0 |
| **Total** | **30** | **82** | **80** |

### Media Preparation

Nutrient broths, Buffered Peptone Water (BPW, CM1049B, Thermo Fisher Scientific) and Tryptic Soy Broth (TSB, 211825, BD Biosciences), were prepared according to the manufacturer’s instructions. Solid culture media, CHROMagar orientation (RT412, CHROMagar), CHROMagar ESBL (ESRT2, CHROMagar), MLGA (CM1031B, Thermo Fisher Scientific), MacConkey no.3 (M7408, Merck) and SCAI (CM0155B, Thermo Scientific, I5125, Merck), were prepared according to the manufacturer’s instructions. Additionally, 1 µg/ml of cefotaxime (sc-202989, Santa Cruz Biotechnology) was added to MacConkey, MLGA and SCAI to provide selection for potential ESBL-producers (22). Agars supplemented with cefotaxime are denoted by ‘+’. A cryoprotectant diluent for -80°C storage of cultures was prepared by dissolving 18.75% w/v maltodextrin (419699, Merck) and 6.25% w/v trehalose dihydrate (90210, Merck) into PBS and filter-sterilised, as developed in (23), with the addition of glycerol to a final concentration of 10%.

### Sample processing and culturing

#### Swab comparison

Foam and polyester swabs were pre-enriched for 18 hours in 5 mL BPW, at 37°C and 220 rpm shaking. After incubation, 10 µL of broth was spread on two agar plates (CHROMagar orientation and CHROMagar ESBL) and incubated at 37°C overnight.

#### Pre-enrichment comparison

Four swabs taken from different areas of the same sampling site (n=82) were pre-enriched in four different conditions: 4 or 18 hours in BPW or TSB, at 37°C and shaking at 220 rpm. After pre-enrichment, the broth was mixed 1:1 with cryoprotectant diluent and stored at -80°C. For use in this evaluation, after approximately six months of storage, the broth was thawed, and an aliquot was diluted 1:100 in PBS, before 50µl was spread onto half of a 2-compartment petri dish containing CHROMagar ESBL and incubated at 37°C overnight. Each petri dish was organised to contain both the 4 and 18-hour enriched broth for the same sample number and broth type.

#### Selective agar evaluation

Swabs were pre-enriched for 18 hours in 5 mL BPW at 37°C and shaking at 220 rpm, the broth was stored 1:1 in cryoprotectant diluent and stored at -80°C. Up to 22 months after storage, 80 samples were selected for use in the agar evaluation. Samples were taken from a collection that had previously been tested for the presence of ESBL-Ec and ESBL-Kp by plating onto both MLGA+ and SCAI+, and species identification using either qPCR or MALDI-TOF MS (Biotyper sirius system, Bruker). From the 80 samples in this evaluation, all grew on cefotaxime-supplemented plates, 39 contained *E. coli* (*23*) and/or *K. pneumoniae* (*18*), and 41 contained only other species (i.e. tested negative for *E. coli* and *K. pneumoniae* previously. The number of samples with confirmed ESBL-Ec or ESBL-Kp was chosen to be approximately half, to challenge our methods with a similar number of positive and negative samples. Broth was diluted 1:100 and 100µl was spread onto MLGA+, CHROMagar ESBL, MacConkey+ and SCAI+ plates and incubated at 37°C overnight.

#### Plating evaluation

The following day, colony colours on the selective agars were assessed for each sample. Interpretation of the colours was based on the manufacturer’s guidance, for our use case ‘positive’ results were limited to our target organisms (*E. coli and K. pneumoniae*). In addition, we noted a high number of non-galactosidase-producing *E. coli* in our samples, so we expanded our interpretation of presumptive *E. coli* colonies on MLGA+ to include yellow, in addition to green and blue colonies. Species were then confirmed using MALDI-TOF MS or qPCR. For identification by MALDI-TOF MS, a single colony was picked with a wooden toothpick and applied to the target plate. Samples were prepared by the extended direct transfer method (Bruker) and tested on the MALDI Biotyper sirius. This method was used for 44% of the pre-enrichment comparison samples and all of the agar comparison samples. For qPCR species confirmation, a duplex qPCR assay was used to determine if suspected colonies were *E. coli* or *K. pneumoniae*. Single colonies were extracted using the MasterPure Complete DNA and RNA Purification Kit (LGC Biosearch Technologies), according to the manufacturer’s instructions. DNA was quantified using the Qubit 4 fluorometer (Agilent Technologies) and standardised to 4 ng/µL. Each reaction consisted of: 10 µL with Luna Universal Probe qPCR Master Mix (New England Biolabs), 0.8 µL forward and reverse primers, 0.4 µL probes, 5 µL of sample DNA and 1 µL of nuclease-free water to a final volume of 20 µL. metH primers and probes were used for *E. coli* detection (Forward: CGTGGTGGTCGCTTTTACCACAGAT, Reverse: TCCACTTTGCTGCTCACACTTGCTC, Probe: [FAM]AAATCTGGGTTRAGCGTGT[BHQ1]), as described in (24), sequences listed in (25)). The recA gene was used for *K. pneumoniae* detection, as recommended in (26), primers and probes were designed in house (Forward: CGCGCACTTTCTTCTCAATC, Reverse: AGCTACAACGGCGACAAA, Probe: [VIC]CGGGTTCTCTTTCAGCCAGGTGAT[BHQ1]. The cycling conditions were: 1 cycle 95°C for 60 seconds followed by 40 cycles of 95°C for 15 seconds, 60°C for 30 seconds. qPCR was performed on QuantStudio 5 and analysed using QuantStudio Design and Analysis software (Applied Biosystems). A Cq threshold was set at 30 cycles. This species confirmation method was applied to the swab comparison samples and 56% of the pre-enrichment comparison samples.

### Statistical analysis

Measurements of bacterial recovery under different conditions were summarised in 2×2 contingency tables, p values calculated with McNemar’s test with continuity correction and odds ratios with 95% confidence intervals using the mcnemar.exact() function in the exact2×2 v1.6.9 package (27) in R v4.4.2 (28). P < 0.05 was considered statistically significant.

## Results

### Comparison of swab types for bacterial recovery

We initially tested two different swab types, foam and polyester, to assess whether one performed better in capturing bacteria from environmental sites within a hospital. We swabbed 30 sites over two hospital wards with both swab types, enriched in BPW for 18 hours and plated onto CHROMagar orientation and ESBL-selective CHROMagar ESBL. From foam swabs, a significantly higher number of samples contained bacteria that grew on CHROMagar orientation, compared to polyester swabs (30 vs 23, p=0.031, OR= 0, 95% CIs= 0-0.849). Although foam swabs also yielded a slightly higher number of samples containing ESBL-producing gram-negatives than polyester swabs (21 vs 15), this difference was not statistically significant (p=0.070, OR= 0.143, 95% CIs= 0.003-1.112) (Figure S1). However, foam swabs were often too large and bulky to sample hard-to-reach areas such as shower drains, without breaking the foam tip. In addition, the foam swabs had a 5.8-fold higher cost compared to polyester swabs (Table S1), we thus chose to continue our sampling with the polyester swabs.

### Pre-enrichment for 18 hours increases recovery of ESBL-producers from environmental swabs

To assess the impact of pre-enrichment conditions on the recovery of ESBL-Ec and ESBL-Kp, we tested samples from 82 sites using four polyester swabs for each site, then pre-enriched in BPW and TSB for 4 and 18 hours (Figure 1).

**Figure 1:**
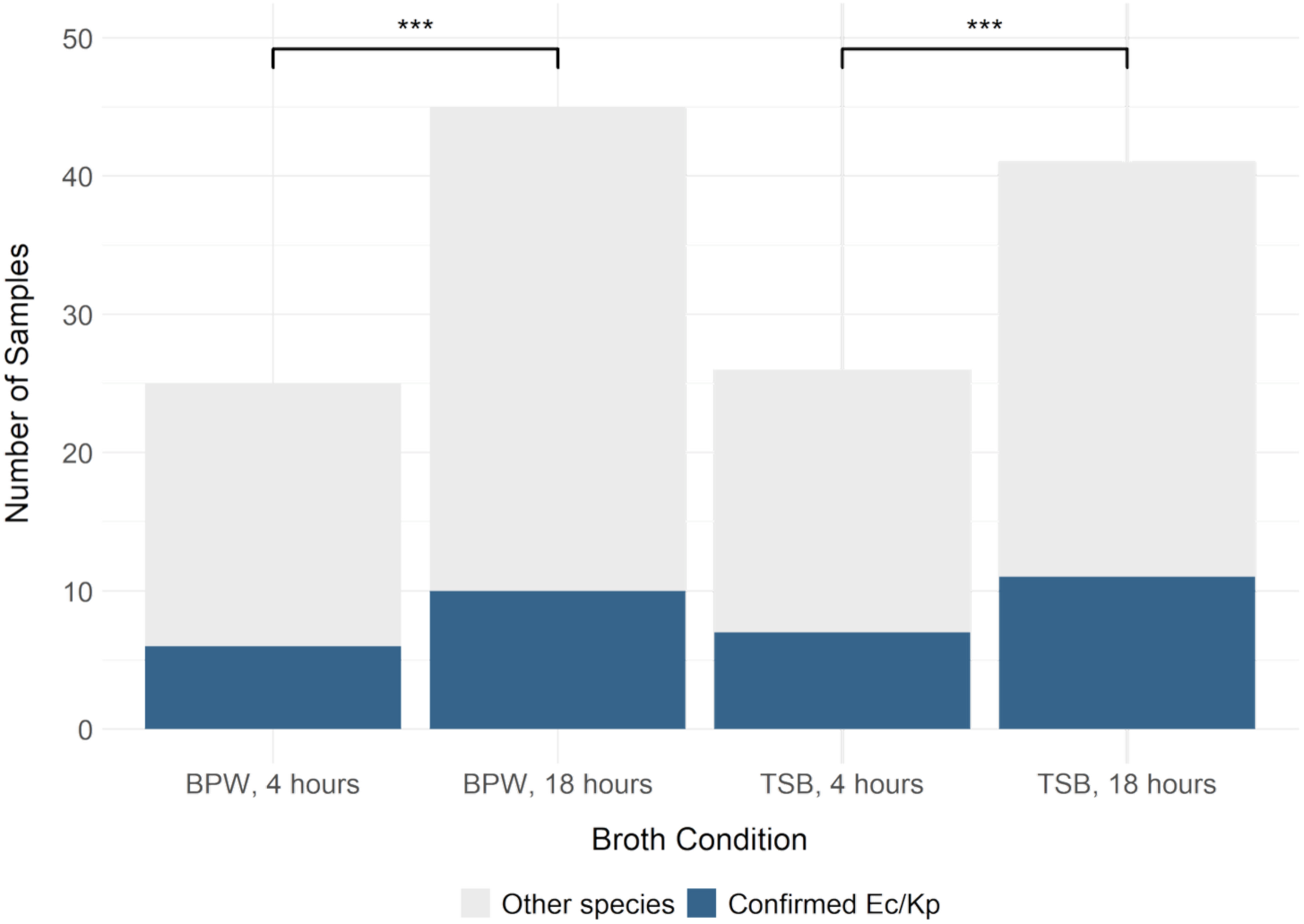
Detection of ESBL-Ec and ESBL-Kp (blue) and other ESBL-producing gram-negatives (grey) from plated environmental samples after pre-enrichment in buffered-peptone water (BPW) or tryptic soy broth (TSB), each incubated for 4 and 18 hours. Significance of pairwise comparisons (McNemar’s test) shown as p<0.001 = ‘***’.

Both enrichment broths performed similarly, and there was no statistically significant difference between the recovery of 3GC-resistant gram-negative-positive samples from BPW and TSB by pairwise comparison at 4 hours (25 vs 26 samples, p=0.724) and at 18 hours (45 vs 41 samples, p=0.387), or for ESBL-Ec and ESBL-Kp (4 hours: p=1.00 OR=0, 95% CIs=0-39, 18 hours: p=1.00 OR=0.667, 95% CIs=0.056-5.82).

Comparison between the two timepoints showed a greater recovery at 18 hours, which was true of pairwise comparisons with each broth. In BPW-enriched samples, 3GC-gram-negatives were recovered from 25 samples at 4 hours, increasing to 45 at 18 hours (p= 5.96 × 10^−8^); with TSB 26 samples were recovered at 4 hours, increasing to 41 at 18 hours (p= 1.91 × 10^−6^). Confirmed ESBL-Ec and ESBL-Kp, numerically increased from 4 to 18 hours (6 to 10 samples for BPW, 7 to 11 samples for TSB) which was not statistically significant (BPW p=0.125, TSB p=0.125, OR=0, CIs=0-1.51). Because of the higher recovery of 3GC-resistant gram-negatives, we decided to enrich our swabs for 18 hours.

The choice of BPW or TSB did not impact the recovery. We continued with pre-enrichment in BPW because of the lower cost of this broth at the time of evaluation (Table S1).

### Differential recovery of ESBL-Ec and ESBL-Kp from different selective agar

We next tested four different selective agars to assess their performance for the detection and recovery of ESBL-Ec and ESBL-Kp from enriched environmental swabs. We plated 80 samples onto MLGA+, CHROMagar ESBL, MacConkey+, and SCAI+, then tested the species identity of each colony of distinct morphology and colour, using MALDI-TOF MS. To ensure our colony selection method would capture the majority of species present on the agar, we first ran a small pilot by plating 2 environmental samples onto each agar, picking 10 colonies of each distinct colony morphology (totalling 120 colonies from 12 morphologies), and testing their species identity using MALDI-TOF MS. All colonies were the same species in 11/12 of morphologies tested, demonstrating that picking a single colony of each colony morphology would not skew the results (Table S2). In addition, to account for instances where multiple colonies with the same identity were picked from one plate, MALDI-TOF MS results were filtered to remove duplicates of the same species.

From the 80 plated samples, colonies with colours indicating ESBL-Ec or ESBL-Kp (ESBL-Ec = yellow/green/blue on MLGA+, pink on CHROMagar ESBL, pink on MacConkey+, N/A on SCAI+; ESBL-Kp = yellow on MLGA+, blue on CHROMagar ESBL, pink on MacConkey+ and yellow on SCAI+) were selected for species confirmation using MALDI-TOF MS. After filtering out any duplicate results, the number of colonies from each plate and target was counted to form the total yield. Results were then split into yield of true ESBL-Ec and ESBL-Kp, and false detection of any other species (Figure 2). To estimate the true number of ESBL-Ec and ESBL-Kp that could be detected in our sub-sample (acting as our gold standard), we calculated the number of samples that were positive on any one plate across all the agars – a putative maximum yield. From this, we could then calculate percentage yield for each agar (Table S3). To test the robustness of this evaluation, we repeated the same experiment using the same 80 samples independently (Figure 2 B). Both experiments were consistent, with the repeat showing slightly higher yield.

**Figure 2:**
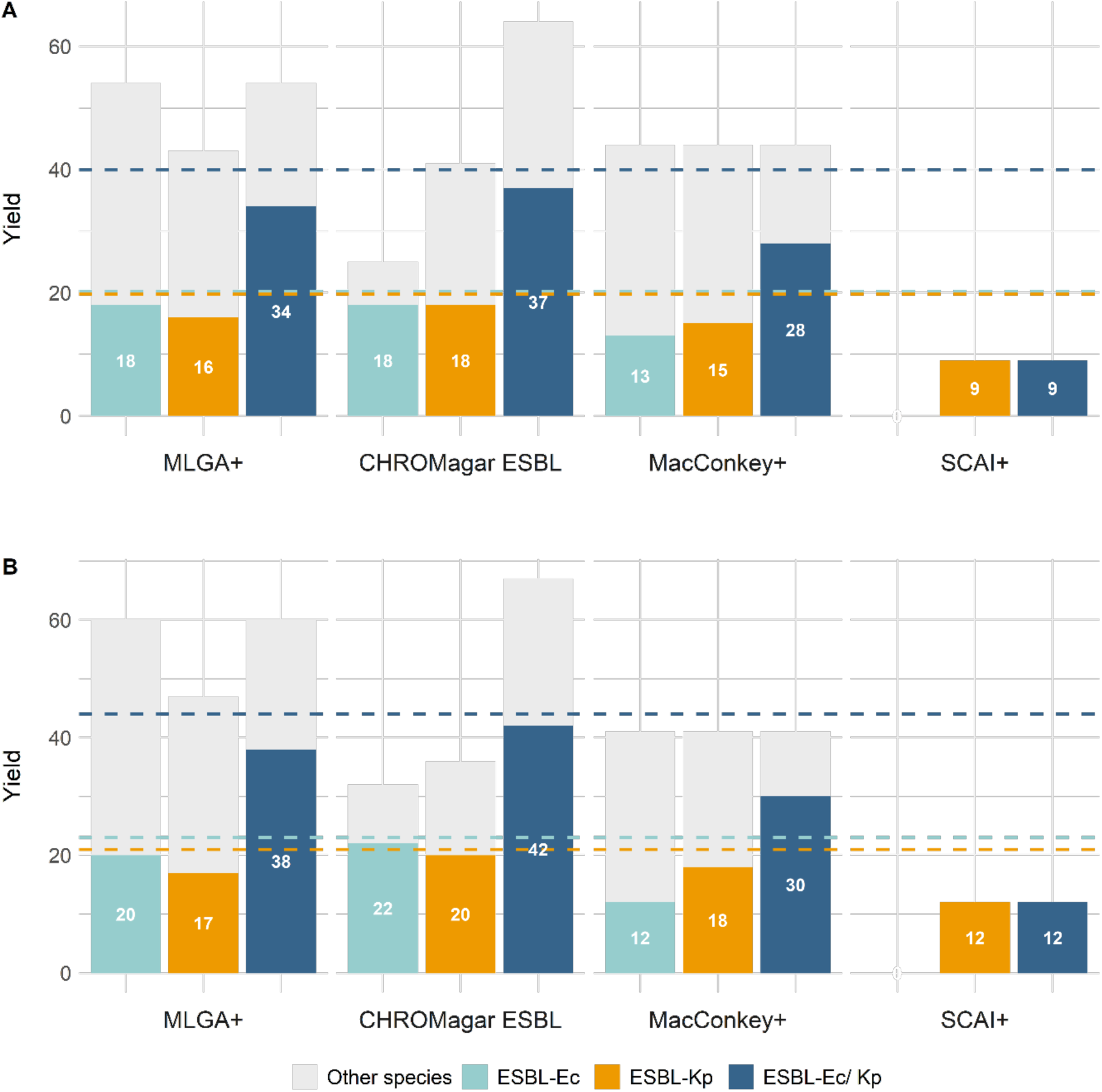
Yield of ESBL-Ec and ESBL-Kp, (ESBL-Ec= turquoise, ESBL-Kp= orange, combined ESBL-Ec and ESBL-Kp= blue) and other 3GC-resistant bacteria (grey) when picking presumed positive colonies from 4 agar types (MLGA+, CHROMagar ESBL, MacConkey+, SCAI+). For each agar, the 3 bars show different targets for the screen: *E. coli*, *K. pneumoniae* and both species combined. The dotted lines represent the true number of ESBL-Ec and ESBL-Kp by combining the results of all agars – a putative maximum yield. The 80 samples were plated and tested for species ID in two independent experiments (A and B).

CHROMagar ESBL demonstrated the best performance for ESBL-Ec screening, demonstrating both a high number of true detections (replicate 1, R1: 18/20 detected, replicate 2, R2: 22/23 detected) and a low number of false detections, with only 7 and 10 (R1 and R2, respectively) confirmed as other species. MLGA+ performed similarly for true detection of ESBL-Ec (R1: 18/20, R2: 20/23) but had higher numbers of falsely detected colonies (36 and 40), due to its lack of differentiation between *E. coli* and *K. pneumoniae*.

For ESBL-Kp screening, true detection was similar in MLGA+ (R1: 16/20, R2: 17/21), CHROMagar ESBL (R1: 18/20, R2: 20/21) and MacConkey (R1: 15/20, R2: 18/21), but all three had a large number of false detections (R1: 27, 23, 29, R2: 30, 16, 23, respectively) as the colour is not specific for *K. pneumoniae* only. On MLGA+ and MacConkey+, all other lactose fermenters present as the same colour as *K. pneumoniae,* and on CHROMagar ESBL *Enterobacter* spp. and *Citrobacter* spp. appear similarly. SCAI+ was the only *K. pneumoniae-*specific agar tested, and the true detection (R1: 9/20, R2: 12/21) was poor, potentially due to a high number of inositol-negative *K. pneumoniae* (18). However, no other species were falsely indicated as ESBL-Kp. Due to the specificity of SCAI to *K. pneumoniae*, we also calculated the yield of ESBL-Kp when using a combination of SCAI+ and one of the three other agars tested. For MLGA+, CHROMagar ESBL and MacConkey+, the addition of SCAI+ increased ESBL-Kp detection between 0-2 isolates, increasing the yield by 0-10% (Table S3).

When combining ESBL-Ec and ESBL-Kp screening, MLGA+ (R1: 34/40, R2: 38/44) and CHROMagar ESBL (R1: 37/40, R2: 42/44) performed similarly overall, with the former detecting fewer ESBL-Ec and ESBL-Kp but benefitting from having fewer false positives (i.e. colonies indicative, but not confirmed, as being *E. coli* or *K. pneumoniae)*. MacConkey+ demonstrated lower detection of ESBL-Ec and ESBL-Kp (R1: 28/40, R2: 30/44) but had a relatively small number of false positives. To favour *K. pneumoniae* growth, SCAI contains carbon sources that are rarely utilised by *E. coli*, and they only appear as ‘tiny, watery colonies’ (18). Therefore, the combined screening result of this agar was the same as when only screening for ESBL-Kp i.e. no ESBL-Ec were detected, as expected.

We found that the optimal choice of agar changed with the species target, whether it was ESBL-Ec, ESBL-Kp, or both species. For detection of both ESBL-Ec and ESBL-Kp, as was the aim for our ongoing surveillance and transmission study (TRACS-Liverpool), CHROMagar ESBL and MLGA+ performed the best. CHROMagar ESBL had better true detection but had more false detection than MLGA+.

### Practical evaluation of selective agar

We next conducted a broader evaluation of the agar, assessing each using both the ESBL-Ec and ESBL-Kp recovery data (Figure 2) and additional practical factors, including cost, ease of preparation (assessed on number of steps and equipment required) and colony characteristics (size and texture of colonies, ease of colour interpretation) that are relevant to their use case in surveillance (Figure 3). A score between 1-10, 10 being the highest performing, was assigned to each category (Table S4).

**Figure 3:**
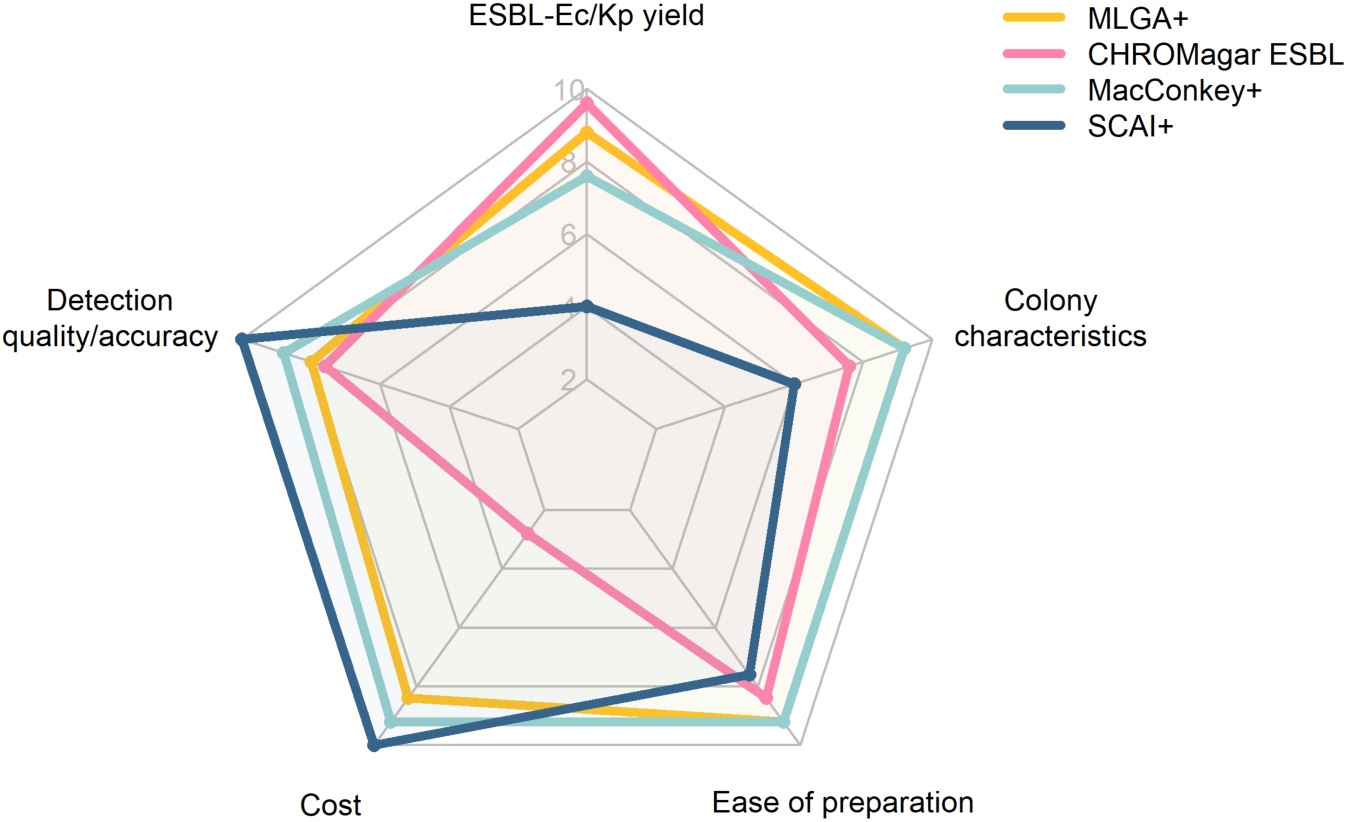
Performance of selective agar scored on ESBL-Ec/Kp yield, detection quality/accuracy, cost, ease of preparation and colony characteristics. Higher scores indicate better performance for each category.

MacConkey+ and MLGA+ achieved the highest total score, and both agars showed a balanced performance, scoring moderately to well in all categories. MLGA+ scored higher for true detection, whilst MacConkey+ scored higher on detection quality and had a lower cost. CHROMagar ESBL performed best for true detection, however, it had low scores for detection quality, ease of preparation, and had the highest cost. SCAI+ performed poorly in true detection, colony characteristics and ease of preparation, but had high detection quality and was the cheapest. To conclude, each agar presents a trade-off between a balanced score between categories, or an excellent score in one category at the detriment of others. Since we were aiming to capture ESBL-Ec and ESBL-Kp from large numbers of samples from care settings for the TRACS-Liverpool study, high scores in all categories were desired, favouring MacConkey+ and MLGA+. The latter captured a high number of ESBL-Ec and ESBL-Kp, we thus chose MLGA+ as our preferred agar for sample screening.

## Discussion

Drug-resistant infections are an increasingly critical threat to public health (29). The accelerating AMR crisis necessitates complex solutions, and infection prevention and control (IPC) plays a key role in preventing infection with potentially untreatable pathogens. IPC interventions are costly; for them to be effective and sustainable, we require an understanding of the transmission pathways of healthcare-associated pathogens. Healthcare settings provide a plethora of opportunities for AMR acquisition and spread (4) and an overlooked but important transmission route is thought to be through environmental exposure. The environment can harbour resistant organisms such as ESBL-E, which can colonise patients and potentially develop into invasive infection (5–7). Thus, environmental sampling is an important aspect of ESBL-E surveillance, aiming to identify endemic transmission links or in outbreak investigations. With limited resources for surveillance, environmental sampling and processing methods must be optimised to be sensitive and cost-effective. Therefore, we have evaluated laboratory methods for screening of environmental samples, comparing the recovery of ESBL-Ec and ESBL-Kp from environmental swabs when using different pre-enrichment broths, pre-enrichment times and selective agar and evaluated practical aspects, including cost.

Our evaluation of enrichment conditions showed higher recovery of 3GC gram-negatives at 18 hours, irrespective of broth type. This increased capture of resistant organisms creates a more comprehensive representation of the microbes in the environment, enabling the capture of more potential transmission events. In contrast, we previously reported that a shorter 4-hour pre-enrichment of stool samples or rectal swabs was superior to a longer enrichment, reducing processing time and the potential for competition bias within the broth, whilst retaining consistent ESBL-Ec detection (30).

Selective agar, although useful to screen for particular organisms in complex samples by inhibiting growth through antibiotic selection and/or metabolic restrictions, has some limitations in identifying the correct species, as metabolic capabilities often overlap between species and can vary within a single species. Our comparison of the yield and accuracy of detection on four agars suggest that performance is dependent on the target species. CHROMagar ESBL had the highest true detection of combined ESBL-Ec and ESBL-Kp, followed closely by MLGA+, which benefited from having fewer false positives. Previously, we found that recovery of ESBL-Ec from pre-enriched stool samples was comparable between CHROMagar ESBL, MacConkey+ and MLGA+ (30). The detection efficacy between agars was more variable in our current study, thus, the performance of these agars is likely dependent on the sample-type.

Our detection assay for ESBL-Ec and ESBL-Kp from agar was robust, and the two experimental replicates showed a similar trend with small variability. The slight increase in yield across R2 is likely explained due to an increase in the number of colonies tested in R2 (R1=399, R2=433). This may be attributed to more conservative identification of distinct morphologies over time, using different batches of agar and small differences between sub-sampling of the pre-enrichment broths during plating.

In addition to our evaluation on detection performance, we considered practical considerations: cost, ease of preparation and growth characteristics of organisms. These are important factors when planning workflows and especially, cost is a limiting factor in many settings. CHROMagar ESBL provided the greatest yield of combined ESBL-Ec and ESBL-Kp and benefits from providing the resolution of differentiating these species, but its cost may be prohibitive.

Our evaluation was limited to a single sample location in Liverpool and only a small number of samples were ESBL-Ec or ESBL-Kp positive. Moreover, we evaluated a limited number of screening conditions, which are not exhaustive of available swab types, broths and agars. We chose these conditions to cover a range of selective agars with different selective properties and intended use-cases, short and long pre-enrichment durations, high-nutrient and recovery nutrient broths.

To conclude, our evaluation provides evidence for the performance of different laboratory screening methods to detect ESBL-Ec and ESBL-Kp from environmental swabs taken from healthcare environments. We found that an 18-hour pre-enrichment in BPW, followed by plating onto MLGA+ worked best overall for our use-case. This provided a high yield of ESBL-Ec and ESBL-Kp to take forward for species confirmation, whilst keeping the workflow practical and low-cost. However, some of the other workflows showed comparable performances, and use of these may be more appropriate in other contexts. Ultimately, this work allows for better informed and context-specific choices when designing laboratory screening workflows for environmental ESBL-Ec and ESBL-Kp surveillance.

## Acknowledgements

We would like to thank the wider TRACS consortium for facilitating this study within the sample collection period. Thank you also to Karina Clerkin for her support in the collection and delivery of the environmental samples, and to the healthcare facility staff for enabling this collection.

## Data availability

All data is available in this manuscript and the supplementary files.

## Supplementary information

Supplementary Figure 1 and supplementary Tables 1-4 are accessible in the supplementary material of this manuscript.

## Author contributions

Conceptualization and method development were done by SF, MM, NAF, JML and FEG.

Investigation was undertaken by EPB, CMK, SF, SG and MM.

Data analysis was done by EPB, CMK, SF and SG.

The original draft was prepared by EPB and FEG.

and then reviewed and edited by all authors.

Supervision was provided by SF, NAF, JML and FEG.

