## Supplementary Figure 1 for "An evaluation of screening methods for the detection of extended-spectrum beta-lactamase-producing *Escherichia coli* and *Klebsiella pneumoniae* in environmental samples from healthcare settings"

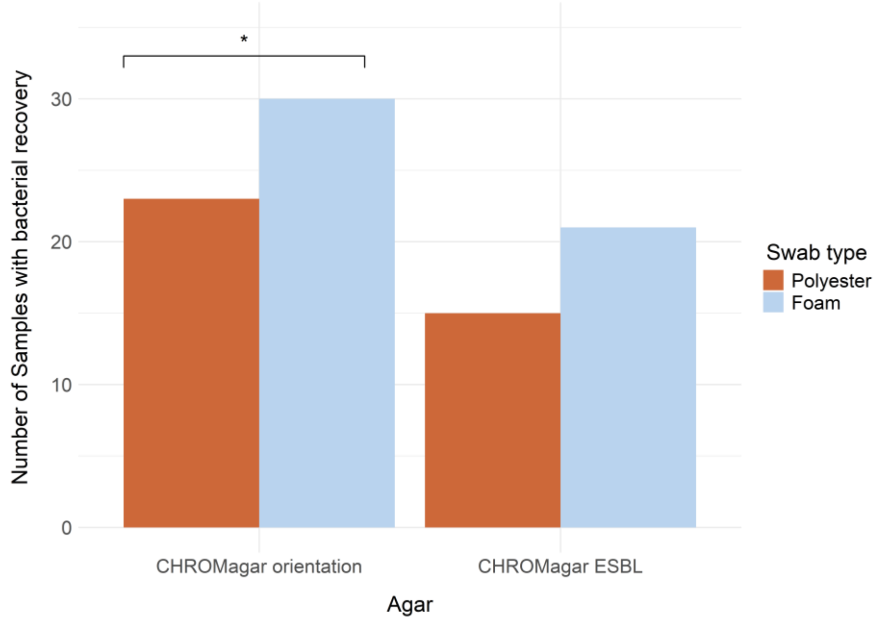


**Figure S1:** The number of samples with growth on CHROMagar orientation and CHROMagar ESBL, selective for ESBL-producing Gram-negative bacteria, when 30 environmental sites were swabbed with 2 swab types- polyester and foam. Significance of pairwise comparisons (McNemar's test) shown as $p<0.05$ = '*'.
